# Effect of buspirone on the H-reflex in acute spinal decerebrated mice

**DOI:** 10.1101/281451

**Authors:** Yann Develle, Hugues Leblond

## Abstract

Pharmacological treatment facilitating locomotor expression also modulates reflex expression through the re-arrangement of spinal networks. Buspirone, a partial serotonin receptor agonist (5-HT_1A_), was recently shown to facilitate and even trigger locomotor movements in mice after complete spinal lesion (Tx). Here, we studied its effect on the H-reflex after acute Tx in adult mice. To avoid possible impacts of anesthetics on H-reflex depression, experiments were performed after decerebration in un-anesthetized mice (N=13). The H-reflex in plantar muscles of the hind-paw was recorded after tibial nerve stimulation 2 h after Tx at the 8^th^ thoracic vertebrae. Average H/M ratio was compared before and every 10 min after buspirone (8 mg/kg, i.p.) for 60 min. Frequency-dependent depression (FDD) of the H-reflex was assessed before and 50 min after buspirone. Before buspirone, a stable H-reflex could be elicited in acute spinal mice and FDD of the H-reflex was observed at 5Hz (68%) and 10Hz (70%) relative to 0.2Hz. After buspirone, the H/M ratio was initially significantly decreased to 69% of pre-treatment. It then increased significantly 30 to 60 min after exposure to buspirone, reaching 170% 60 min after injection. FDD 50 min after buspirone was not significantly different that FDD without treatment. Altogether results suggest that the reported pro-locomotor effect of buspirone occurs at a time where there is a reflex depression followed by a second phase marked by enhancement of reflex excitability, denoting functional inhibitory control.

**KEY POINTS SUMMARY:**

- Buspirone, a partial serotoninergic agonist (5HT_1A_), exerts a considerable acute facilitation of spinally-mediated locomotion after a complete spinal lesion in mice.
- Since locomotion is associated with reflex modulation, we focused in this study on the effect of buspirone on H-reflex in acute spinal mice after decerebration and removal of anesthesia.
- Buspirone injection resulted in a depression of the H-reflex during the first 20 minutes followed, 40 minutes later, by an increase of the reflex.
- The H-reflex increase is not due to a disinhibition since buspirone had no effect on the frequency dependent depression (FDD) of the H-reflex.
- The use of a model of adult decerebrate mouse with complete spinal cord injury allows to establish that buspirone has a biphasic effect on the H-reflex and that the inhibitory action on sensory information concurs with an excitatory action on locomotion.

**Abbreviations:**

5-HT: 5-Hydroxytryptamine
5-HT_1A_: 1A subunit in 5-Hydroxytryptamine receptor
8-OHDPAT: *8*-Hydroxy-2-DI-n-Propylamino-Tetraline CPG Central Pattern Generator
EMG: electromyographic
FDD: Frequency Dependent Depression
GABA: Gamma-Aminobutyric-Acid TASK-1 acid-sensitiv-K-1
Tx: complete transection

## INTRODUCTION

During locomotion, afferent inputs from the hind limbs serve to control the excitability of spinal networks. They adjust motor output by direct impact on either motoneurons or interneurons, comprising those of the central pattern generator (CPG) that is responsible for locomotion (Rossignol,2006). After complete spinal cord injury, sensory feedback becomes the only source of input remaining to the spinal cord: it has the power to re-arrange spinal circuits below the lesion, as shown by the positive outcome of treadmill training in adult cats (Barbeau and Rossignol,1987, Belanger et al.,1996, Lovely et al.,1986), rats (Edgerton et al.,1997, Ichiyama et al.,2008, Otoshi et al.,2009) and mice (Leblond et al.,2003). The plasticity involved in this recovery of locomotion necessarily entails changes in several reflex pathways (Côté and Gossard,2004, Côté et al.,2003).

In spinal animals, pharmacological treatments that mimic neurotransmitters from severed, descending fibers also have neuromodulator effects on locomotor networks and can improve recovery of locomotion (Chau et al.,1998a, b). As is the case with locomotor training, drugs that enable functional recovery also regulate spinal reflexes (Côté et al.,2003, Frigon et al.,2012). For example, in cats with complete spinal lesion, the noradrenergic agonist clonidine, which is known to trigger hind limb locomotion (Barbeau and Rossignol,1987), was also found to modify spinal neuron responses to peripheral inputs (Barbeau and Rossignol,1987, Chau et al.,1998a, Côté et al.,2003, Frigon et al.,2012). In rodents, serotoninergic (5-HT) drugs are effective in triggering and facilitating locomotion after complete spinal lesion (Schmidt and Jordan,2000, Slawinska et al.,2014). Recent work in our laboratory has established that treatment with the US Food and Drug Administration-approved 5-HT_1A_ receptor partial agonist buspirone (Loane and Politis,2012) can initiate locomotion in the hind limbs of adult mice immediately after complete spinal lesion (Jeffrey-Gauthier et al.,2017). As drugs with pro-locomotor properties also modify reflex pathways, buspirone may alter reflex excitability in mice after complete spinal lesion.

The effects of 5-HT_1A_ agonists on spinal reflexes have been tested earlier in different animal models, but there is still no consensus today as to whether the outcome is excitatory or inhibitory. On the one hand, *in vitro* results on isolated brainstem and spinal cord in neonatal rats indicate that buspirone decreases monosynaptic reflex excitability (Yomono et al.,1992). This observation concurs with other studies that have demonstrated 5-HT_1A_ receptor inhibition in reflex pathways (Crick et al.,1994, Hasegawa and Ono,1996a, Hasegawa and Ono,1996b, Honda and Ono,1999, Nagano et al.,1988). On the other hand, some have reported excitatory effects of 5-HT_1A_ (Clarke et al.,1996), mainly by showing facilitatory effects on motoneuron depolarization (Grunnet et al.,2004, Perrier et al.,2003, Takahashi and Berger,1990, Zhang,1991) or monosynaptic reflex enhancement (Honda and Ono,1999). Is it possible that substances with excitatory effects on locomotion also have inhibitory effects on spinal cord excitability?

The present study was performed with a newly-developed model of decerebrated mice and was designed to investigate the modulation of reflex pathways in the absence of pharmacological anesthesia. This was required, since locomotion involves wide re-organization of reflex pathways, as shown mainly in decerebrated cat preparations in which new relays were described in the absence of anesthesia (McCrea,2001). Some reflex pathways are thus state-dependent, meaning that they occur only when the CPG is driving locomotion or when drugs known to trigger locomotion are given (Gossard et al.,1994, Leblond et al.,2000, Leblond et al.,2001, Perreault et al.,1995).

Here, the main objective is to assess the effect of buspirone, at a dose level that triggers locomotion, on H-reflex amplitude in adult decerebrated mice after acute spinal cord lesion. This reflex, the electrical analog of the tendon tap reflex, is primarily mediated by monosynaptic pathways (Misiaszek,2003). The results show a biphasic effect of buspirone on the H-reflex: its amplitude is first decreased significantly, then increased significantly, starting 30 min after drug administration. This increase is still evident 60 min after buspirone and is not associated with any significant changes in frequency-dependent depression (FDD) of the H-reflex. Some of these results have been presented in abstract form (Develle and Leblond,2016).

## METHODS

### Animal care and ethics

Experiments were performed on 13 C57 mice, of either sex (Charles River Laboratories, Saint-Constant, Québec, Canada), weighing 20 to 30g. Their living conditions were strictly controlled by laboratory and facility staff. They were housed in cages with food and water available *ad libitum*. All manipulations and procedures were in accordance with Canadian Council on Animal Care guidelines and were approved by the Université du Québec à Trois-Rivières Animal Care Committee. The mice were randomly assigned to 1 of 2 groups in acute, terminal experiments to evaluate the effect of buspirone on the H-reflex: a test group (n=8) exposed to buspirone and controls (n=5) treated with saline.

### Anesthesia

All surgeries were performed under isoflurane anesthesia (2% mixed with O_2_ 95% and CO_2_ 5%, 200 ml/min). General anesthesia was first induced through a mask: then, the animals were tracheotomized to maintain anesthesia and allow artificial ventilation (SAR-830/P Ventilator, CWE, Inc., Ardmore, PA, USA) adjusted to preserve expired CO_2_ level between 3 and 4% (CapStar-100 CO_2_ monitor, CWE, Inc.). Body temperature was monitored by rectal probe and maintained at 37 ±0.5°C with heating pad.

### Spinalization

The objective was to measure the H-reflex after complete spinal cord section. It was performed early in the surgery to minimize the impact of the decerebration on the spinal circuitry. The paravertebral muscles were cleared from both vertebral laminae after skin incision targeting the 8^th^thoracic vertebra. Then, double laminectomy exposed the spinal cord at this level. After perforation of the dura mater with a needle, a small piece of lidocaine-soaked cotton (xylocaine 2%) was applied for 1min to prevent uncontrolled secondary neural damage or lumbar spinal cord excitotoxicity. Then, the spinal cord was transected with micro-scissors and confirmed by visual observation of the gap between the rostral and caudal stumps. Finally, Surgicel^®^ absorbable hemostat (Ethicon, Johnson & Johnson, USA) was inserted between the 2 parts of the spinal cord before the skin was sutured.

### Decerebration

Spinal network activities are traditionally assessed in decerebrate preparations, especially from cats, rats and rabbits, with recent adaptation to mice (Dobson and Harris,2012, Meehan et al.,2012, Meehan et al.,2017). Data were, therefore, acquired in decerebrated, unanesthetized mice to avoid the unwanted effects of anesthesia. The carotid arteries were first ligated to minimize cerebral perfusion while the animals were secured in a stereotaxic frame (Model 980 Small Animal Spinal Unit, Kopf Instruments, Tujunga, CA,USA) equipped with a small mouse and neonatal rat adaptor (StoeltingCompany, Wood Dale, IL, USA). They were then craniotomized, taking care to leave the superior sagittal sinus intact. Bone wax (Ethicon, Johnson & Johnson, USA) was applied to the skull when necessary to prevent bleeding. The Dura mater was removed gently to expose the cortex for transection with a razor blade 1mm rostral to the lambda. The rostral part of neural tissue and the occipital cortex were removed, by gentle suctioning with an adapted micro-vacuum, corresponding to pre-collicular-pre-mamillar decerebration. The cavity was finally filled with Gelfoam^®^ thin soak hemostat sponge (Pfizer Inc, New York, NY, USA), and the skin was closed with suture.

### H-reflex recording

After decerebration, the left hind limb was fixed in extension and an incision was made on top of the gastrocnemius muscles to separate and expose the tibial nerve. A pool was formed with skin flaps and filled with mineral oil to avoid nerve desiccation. The tibial nerve was mounted on a home-made bipolar hook electrode for stimulation. One-ms single-pulsed stimulations were delivered by a constant-current stimulator (Model DS4, Digitimer Ltd., Welwyn Garden City, UK) triggered by a computer-controlled sequencer (Power 1401 acquisition system, Cambridge Electronic Design, Cambridge, UK).

Paired, fine, multistrained stainless steel wires (AS631Cooner Wire, Chatsworth, CA, USA) were inserted under the skin, between the 2^nd^and 3^rd^medial toes, towards the intrinsic foot muscles, for electromyographic (EMG) recording. Signals were amplified 1,000x, bandpass-filtered at 30-3,000Hz (Grass P55 AC Preamplifier, Natus Neurology, Inc., Pleasanton, CAUSA), and digitized for data acquisition (Spike 2 software, Cambridge Electronic Design, Cambridge, UK). A ground electrode was inserted in the skin between the stimulating and recording electrodes.

Anesthesia was stopped, followed by 60-min rest, which corresponds to approximately 120-150 min post-spinalization, to avoid undesirable anesthesia-induced effects.

### Buspirone administration

The H-reflex was compared in groups of mice that received buspirone or saline. A catheter was inserted to facilitate i.p. administration without moving the animals. Buspirone (8mg/kg,i.p.) was given in a volume of 0.1cc, with the controls receiving the same amount of saline (0.9%).

### Data acquisition and analysis

Stimulus-response curves (e.g. Fig. 1B) were charted by gradually increasing tibial nerve stimulation intensity to ascertain the maximal H-reflex (4-6 ms latency) concomitant with stable M-wave (1-3 ms latency). At this intensity, which corresponded to approximately 1.8 times the motor threshold, Ia muscle spindle afferents were mainly activated. Responses to tibial nerve stimulation intensity were recorded before and every 10 min after the injection, for a total of 60 min. It allowed us to observe the evolution of reflex amplitude with time and treatment.

**Figure 1.**
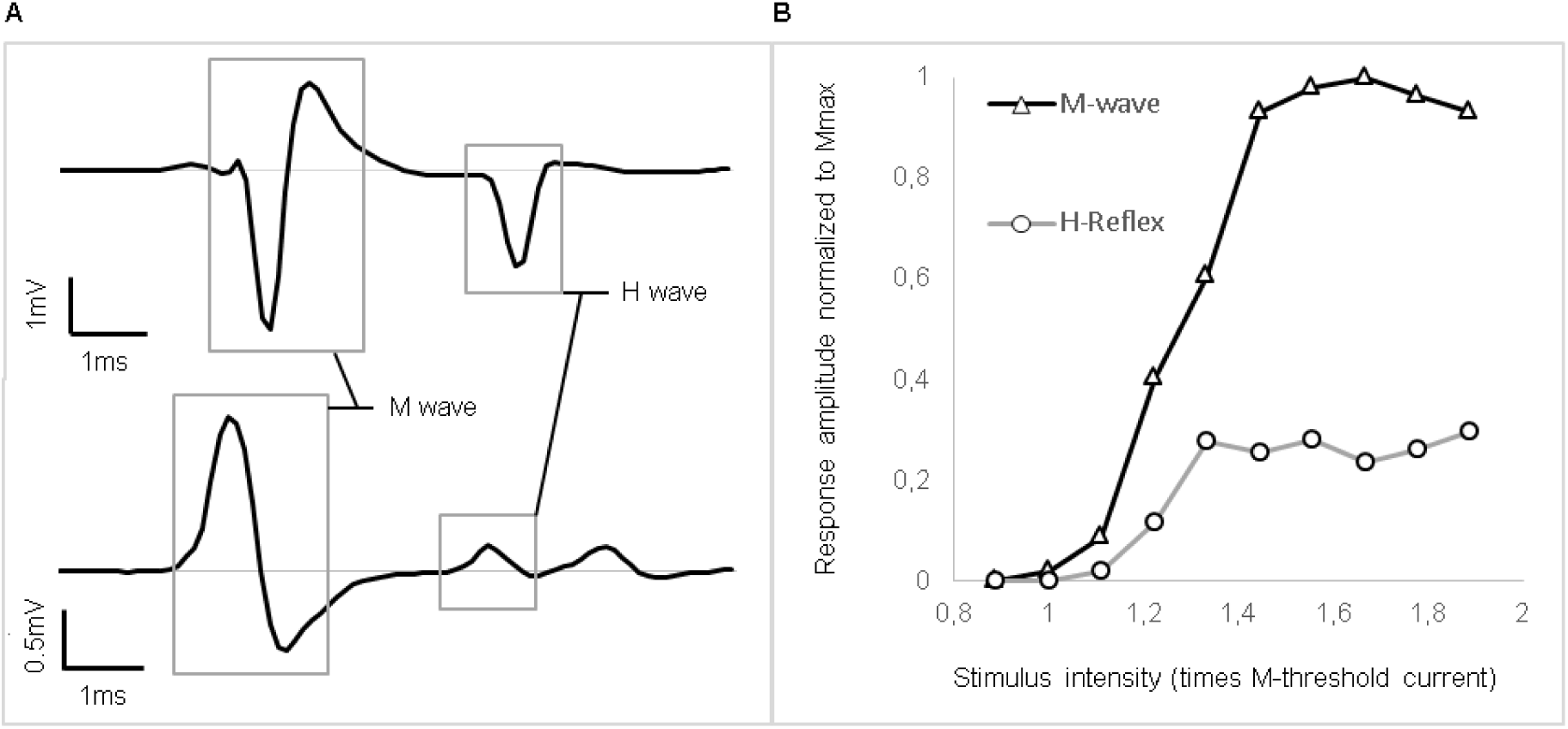
Raw traces of recorded signals and representative recruitment curve. **A)** EMG examples of raw traces recorded. The M-wave is the depolarization of the whole motoneuron pool activated by stimulation, whereas the H-wave is the motor response induced by primary afferent depolarization. **B)** Peak-to-peak amplitude of EMG responses recorded in intrinsic foot muscles by progressively increasingtibial nerve stimulation intensity. This stimulus-response curve is tested to find the stimulation intensity that will allow reflex amplitude evaluation. It should be around Hmax, which specifically activates Ia primary afferents, but stable M-wave response is also necessary to ensure that the stimulation electrode is stable, at the beginning of the M-wave plateau. In this particular example, stimulation around 1.4 times the motor threshold would be chosen.

Spinal inhibitory processes were also tested before and 60 min after injection by varying stimulation frequency between 4 blocks of 30 stimulations (0.2, 5, 10 and 0.2 Hz; 60s inter-block interval) to assess FDD. The first 5 responses of each block were discarded to allow H-reflex stabilization. Analyses comprised only recordings with stable M-wave throughout the protocol (<10% variation) to ensure recording stability. Note that, as illustrated in Figure 1A (bottom trace), some responses included a third deflection with longer latency beginning about 7-8 ms post-stimulation. This response was poorly depressed by stimulation frequency, in contrast to the response localized between 4 to 6 ms, and was not taken into account in H-reflex measurement.

Data were analyzed with Spike2 software (Cambridge Electronic Design) and Excel software (Microsoft Corporation, Redmond, WA, USA). Peak-to-peak amplitudes of the H-reflex and M-wave were measured to establish the H/M ratio so that the results could be compared between animals. Mean ratio at each time point was computed by averaging 30 stimulations at 0.2Hz. In the FDD protocol, mean H/M ratio was averaged from 25 successive responses for each block.

Statistical analysis was conducted with Statistica software (version 13, Statsoft Inc., Tulsa, OK, USA), and the significance threshold was set at p ≤ 0.05. Normal distribution was assessed by the Kolmogorov-Smirnov test. Then, Greenhouse-Geisser-corrected 2-way ANOVA ascertained the effects of the intra-subject factor *time* and the inter-subject factor *treatment* on reflex amplitude. *Post hoc* Student’s t-test targeted periods presenting significant variations in comparison to pre-injection values. Finally, FDD at 60 min post-injection in buspirone-injected animals was tested by 2-way ANOVA to compare with inhibition in the pre-injection state.

## RESULTS

Buspirone administration decreased spinal cord excitability in the first 20 min, followed by its increase 40 min later. This finding was indicated by reduction of H-reflex peak-to-peak amplitude early after treatment, followed by augmentation 40 min later.

### H-reflex in acute spinal decerebrated mice

Stimulus-response curves were recorded for each mouse to establish at which intensity the H-reflex should be evoked to test the effect of buspirone. Typical examples of the H-reflex and stimulus-response curves are depicted in Figures 1A and 1B, respectively. Stimulation intensity was increased progressively until the whole pool of fibers in the tibial nerve was recruited, as indicated by a plateau being reached in the M-wave in Figure1B. The H-reflex was usually evoked close to the motor threshold, and after an initial rise, it too plateaued and did not manifest a classical decrease after reaching maximum. This pattern was observed in all animals (N=13). Stimulation intensity was selected so that stable M-wave could be evoked as near as possible to beginning of the plateau (1.4T in the example depicted in Figure 1B).

### Buspirone effect on the H-reflex

Figure 2A displays H-reflex raw traces averaged from 30 stimulations recorded from intrinsic foot muscles before, 10 and 40 min after buspirone administration in 1 mouse. Average H-reflex decreased dramatically in this mouse 10 min after buspirone (grey trace), then increased considerably 40 min later (dotted trace).

**Figure 2.**
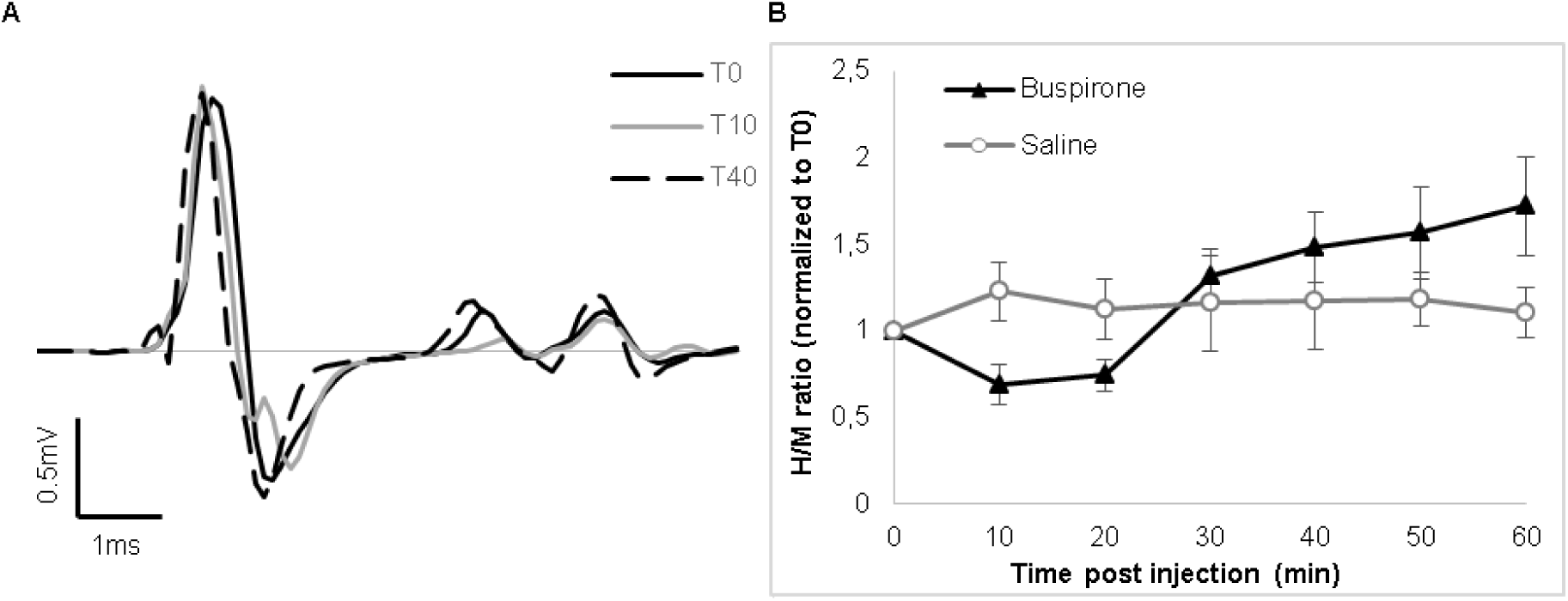
Effect of buspirone on the H-reflex. **A)** Mean traces obtained at 3 critical timepoints by averaging 30 stimulations at constant intensity in abuspirone-treated mouse. Peak-to-peak measurement of H-reflex amplitude shows an initial decrease at T10 (grey trace) *vs* T0 (black trace) and an increaseat T40 (dotted trace). **B)** Representation of reflex amplitude evolution normalized to the pre-dose value by averaging all mice from both groups. Reflex amplitude is significantly decreased during the first 20 min post-dose and significantly increased after 40 min in buspirone-treated animals but not in saline-injected controls.

Figure 2B illustrates that the H/M ratio (normalized to T0) changes over time in the buspirone but not in the saline group. In all groups combined, Greenhouse-Geisser-corrected 2-way ANOVA of the factors time and treatment indicated reflex amplitude variation over time (main effect: F_6,64_= 2.71,p = 0.020). Moreover, this effect of time was associated with the *treatment* applied (F_6,64_ = 3.61, p= 0.004). Indeed, *posthoc* Student t-test first shows reflex amplitude being decreased by 31% by buspirone at T10, significantly different from T0 values (p= 0.012). This reflex inhibition was still significantly different at T20 (p= 0.040). Then, a transition phase led to a significant 48% increase of the H/M ratio from 40 min (p= 0.009) that plateaued at 60 min (T0-T50, p= 0.036, and T0-T60, p= 0.031). In contrast, *post hoc* test ingrevealed no significant differences in the saline-injected group over time (p > 0.05).

### FDD of the H-reflex

The H-reflex was characterized by frequency-dependent behavior. As stimulation frequency was increased from 0.2to 10Hz, reflex amplitude was depressed in intact and acute spinal animals. Figure 3A is a typical example of FDD of the H-reflex in a mouse before buspirone treatment: higher frequency of stimulation at 5 or 10 Hz almost completely abolished the H-reflex. Figure 3B shows inhibition at 5 and 10Hz in all buspirone-treated animals as reflex amplitude normalized to values obtained at 0.2Hz frequency. Our results reflect significant reflex inhibition, depending on stimulation frequency (2-way ANOVA: F_1,36_= 438, p<0.001). Modulation of recorded reflex amplitude tended to be increased by treatment at 60 min post-dose (66.6% at 5Hz and 68.9% at 10Hz at pre-dose versus 78.6% at 5Hz and 75.8% at 10Hz 60 min post-dose). However, the difference was not significant, since statistical analysis did not detect any effect of treatment on the amplitude of frequency-dependent inhibition (2-way ANOVA: F_1,36_ =0.73, p=0.408).

**Figure 3.**
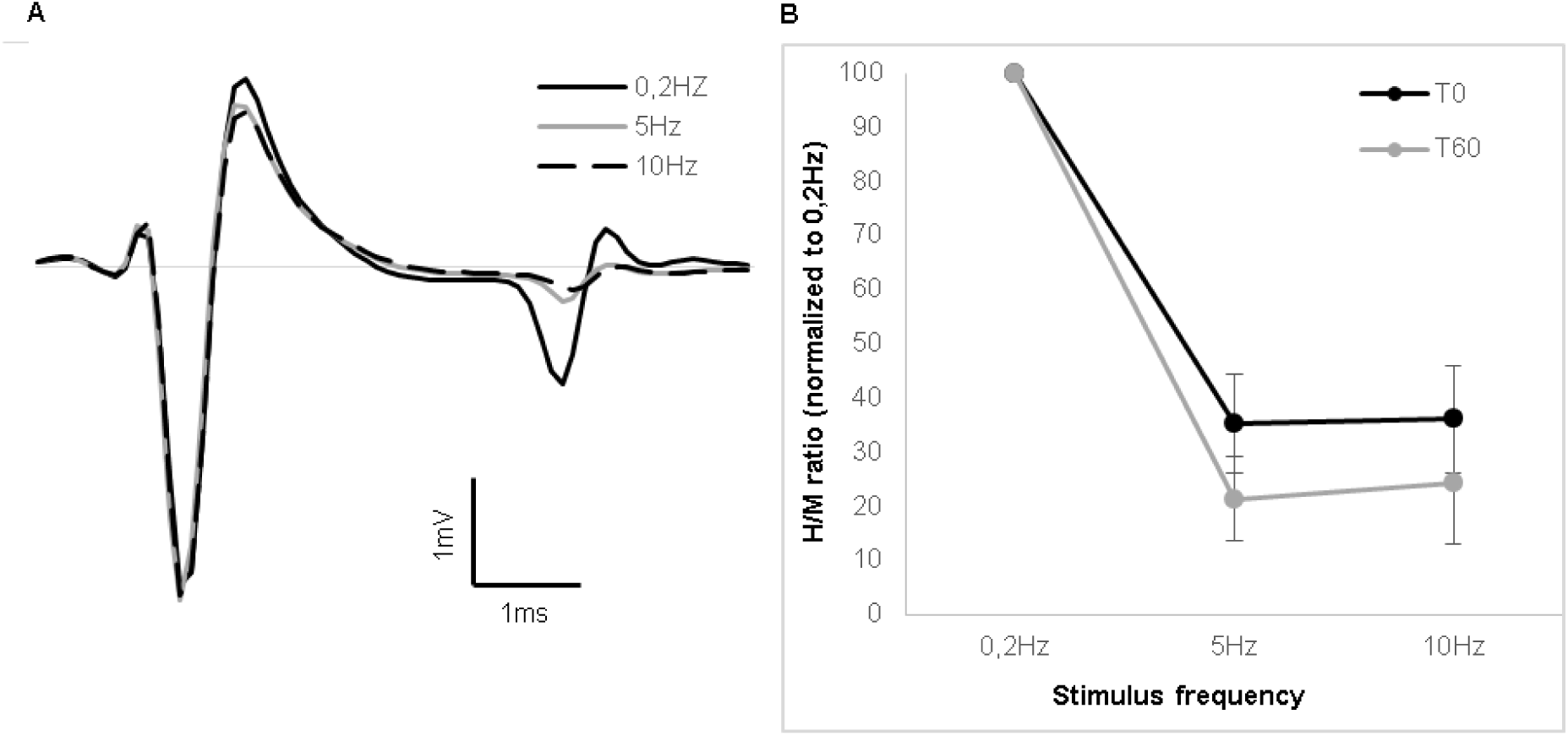
Reflex inhibition at 5Hz and 10Hz. **A)** Mean traces of 25 stimulations at 0.2 Hz, 5 Hz and 10 Hz in a mouse at pre-dose. Consistency in fiber recruitment by stimulation is assessed by stable M-wave between each trial. Only the H-reflex is depressed by stimulation frequency. **B)** Representation of mean reflex amplitude at different stimulation frequencies normalized to values at 0.2 Hz for the whole group before and 60 min after buspirone administration. Inhibition at 5 and 10 Hz does not present any significant difference between the 2 time points.

## DISCUSSION

The use of adult decerebrated mouse preparations allowed us to study the effect of buspirone on the H-reflex after acute spinal lesion in a system that was not altered by the presence of anesthetic drugs. The main study result was that buspirone had a depressive impact on H-reflex amplitude for the first 20 min after drug administration. This depressive outcome was then attenuated and even reversed to an increased effect on the H-reflex which became significant 40 min post-dose. Since FDD at 5 and 10 Hz remained similar to the pre-dose condition, the reflex enhancement observed at 40 min did not seem to be attributed to loss of inhibitor control.

### Buspirone act as a 5HT_1A_receptor agonist on the reflex

Buspirone is a selective partial agonist of 5-HT_1A_ receptors with some affinity for dopaminergic D2 receptors (Loane and Politis,2012). Since it has been shown that motoneuronal excitability is not affected by either D2 agonist or antagonist (Han et al.,2007, Han and Whelan,2009), we assume that the effect observed in our experiment on the H-reflex with buspirone is mainly associated with 5-HT_1A_ receptors (see also D’Amico *et al*, 2017). Even if this reflex is mainly of monosynaptic nature, it remains under the control of different elements (Misiaszek,2003). The 5-HT_1A_ receptors can be found at various locations on these elements including presynaptic, intrasynaptic and even outside the synaptic innervation. This heterogeneity probably explain why 5-HT have such multiple and opposite effects on motor circuits of the spinal cord, as recently elegantly reviewed in Perrier and Cotel (2015).

The short-term impact of buspirone observed in our experiments, i.e reflex reduction, concurs with the literature on the effect of 5-HT_1A_ agonists. Indeed, monosynaptic reflex reduction has been shown with the 5-HT_1A_ agonist 8-OHDPAT in rats with complete spinal lesion under α-chloralose or urethane anesthesia (Hasegawa and Ono,1996a, Honda and Ono,1999, Nagano et al.,1988). More recently, (D’Amico et al., 2017) used buspirone as 5HT_1A_ agonist and showed that its systemic administration in awake humans reduces about 30% of F-wave amplitude, indicating direct decrease in motoneuron excitability and output.

Because 5-HT_1A_ receptors are mainly present on dorsal laminae of the spinal cord, it was proposed to be also involved in afferent regulation (Giroux et al.,1999, Noga et al.,2009). Such participation in afferent modulation by 5-HT_1A_ receptors has been confirmed by (Hasegawa and Ono,1996b) in rats. They showed that monosynaptic reflexes evoked by dorsal root stimulation are depressed by 8-OHDPAT administration with no change in motoneuronal excitability. This suggests that reflex depression is induced by lowering neurotransmitter release at the presynaptic level. Such afferent regulation could be generated by 5-HT_1A_ receptors on afferent neurons and could be responsible for increased GABA-mediated inhibition (Gharagozloo et al.,1990).

### The absence of anesthesia and reversal from inhibitory to excitatory effects of buspirone

A reversal of the effect buspirone (or any other 5HT_1A_ agonist) from inhibitory to excitatory later post-treatment has not been reported so far. Such differences with previous experiments could be related to the use of decerebrated preparations and the absence of anesthesia (Meehan et al.,2017) that affect reflex modulation (Ho and Waite,2002) see also (Schmidt and Jordan,2000). Indeed, for example, experiments on decerebrated cats disclosed reversal of group I autogenetic inhibition to polysynaptic excitation in extensor motoneurons after exposure to clonidine or L-DOPA, drugs that promote locomotion in spinal cats (Conway et al.,1987, Gossard et al.,1994). Interneurons involved in this reflex reversal are shared with CPGs and supraspinal inputs (Leblond et al.,2000, Leblond et al.,2001).

Low threshold stimulation like the one used in the present study might also activate oligosynaptic pathways by some other large-diameter afferent fibers, such as type Ib afferents, that are also in contact with motoneurons. Because such depolarization involves the polysynaptic circuitry, the motor response would have longer delay and may be dissociated from monosynaptic activation (e.g. Figure 1A, bottom trace). Still, long-lasting effects on motoneurons by these pathways are not to be excluded and could be implicated in signal amplitude recorded by EMG.

Indeed, the absence of anesthesia most likely allowed otherwise quiescent spinal networks to be active and participate in the modulation of membrane conductance, affecting motoneuronal responsiveness (Harvey et al.,2006a, b, Li et al.,2006, Murray et al.,2010) see also (D’Amico et al.,2014). However, slowly-activated currents, like persistent inward currents, required long-duration input and could not be fully actuated by brief stimulations like the ones used in our study (Murray et al.,2011).

It is still not fully clear why the H-reflex was enhanced during the second phase of our experiment and this will be discussed in a later section. Nonetheless, to evaluate if this reversal from inhibition to excitation can be explained by a disinhibition, we measured FDD of the H-reflex. This inhibition is reported to mainly depend on GABA inhibitor potential (Jolivalt et al.,2008, Kakinohana et al.,2006). The results showed that post-activation inhibitory circuitry remained unchanged between pre-dose and 60 min post-dose in our experiment, suggesting that disinhibition or changes in GABAergic activity cannot explain the observed increase of reflex amplitude.

### Opposite effect of 5HT1A according to receptor location on the motoneuron

As mentioned above, 5HT_1A_ receptors are located at various locations that can get activated simultaneously. On the one hand, at the synaptic level, 5-HT is responsible for modulating fast-activated potassium channels via 5-HT_1A_ receptors (Jackson and White,1990, Penington and Kelly,1990). It was shown that 5HT_1A_ receptors inhibits TASK-1 potassium channels that would contribute to the excitatory effect of 5-HT on spinal motoneurons (Perrier et al.,2003). By lowering outward cation flux, 5-HT_1A_ receptors shortened the refractory period and facilitated motoneuronal depolarization (Grunnet et al.,2004, Santini and Porter,2010). This mechanism augments motoneuronal excitability and enhances motor responses to synaptic stimulation. On the other hand, there are extrasynaptic5-HT_1A_ receptors that can be activated by spill-over during high 5-HT release at the synaptic level or background concentration of 5-HT (e.g. in systemic administration) that are known to be inhibitory. Indeed, 5-HT_1A_ receptor stimulation on axon hillocks elicits inhibition of sodium channels that are responsible for initiation of action potentials in motoneurons (Cotel et al.,2013, Perrier and Cotel,2015, Perrier et al.,2013, Petersen et al.,2016). This inhibition decreases the number of spikes triggered and consequently reduce the amplitude of the EMG.

Thus, when large dose of buspirone is given, as in our experiments, 5-HT_1A_ receptors inhibit motoneuron output and decrease reflex amplitude through activity at the axon hillock sites even if there is an excitation at the synaptic level. This dual effect of 5-HT_1A_ receptors on motoneuronal excitability may be involved in the observed biphasic effect of buspirone over time on reflex amplitude through a switch in dominance of receptor type activity.

Indeed, drug action is concentration-dependent, and buspirone pharmacokinetics undergoes a biphasic elimination cycle (Sethy and Francis,1988). The first half-life of the drug is reached after 24.8 min, a period that matches the transition phase of reflex amplitude in treated animals. Also, synaptosomes, the main structures implicated in 5-HT re-uptake in synapses (Henn and Hamberger,1971), are not present at the axon hillock level. This region relies mainly on the participation of astrocytes that have been demonstrated to be involved in 5-HT re-uptake, especially at the extrasynaptic level (De-Miguel et al.,2015, Henn and Hamberger,1971, Kimelberg and Katz,1985, Ritchie et al.,1981). Such region-dependent differences in the 5-HT clearance mechanisms could explain the biphasic effect of buspirone on reflex amplitude over time. Moreover, a desensitization of 5-HT_1A_ receptors after their pharmacological activation have been reported and should be considered as well in that reversal (Seth et al.,1997).

### Reflex inhibition concomitant with excitatory effect on locomotion

Despite its initial inhibitory effect on the H-reflex shown in the present study, it was shown that buspirone promotes locomotor movements after spinal cord injury. Indeed, in another experiment from our laboratory it was shown that buspirone exerts a considerable acute facilitation of spinally-mediated locomotion in mice after a complete section of the spinal cord (Jeffrey-Gauthier et al.,2017). Buspirone was also shown to potentiate locomotion when combined with other treatments (Gerasimenko et al.,2015, Ung et al.,2012). It indicates that locomotion can be triggered during depression of sensorimotor excitability induced by buspirone. This paradox is also observed with the 5-HT_1A_ partial agonist 8-OHDPAT, which is known to inhibit the monosynaptic reflex (Hasegawa and Ono,1996a, Honda and Ono,1999, Nagano et al.,1988) and can facilitate recovery of locomotor function in spinal rats (Antri et al.,2005, Antri et al.,2003). Sensory inputs provided by the treadmill seem sufficient to initiate and maintain locomotor rhythm with buspirone. The same observation was made in cats where clonidine, a noradrenergic agonist that can trigger locomotion on a treadmill after a complete spinal lesion, reduce reflexes evoked by stimulation of the dorsum of the foot (Barbeau and Rossignol,1987, Chau et al.,1998a).

These observations with adult animals that walk on a treadmill seem to disagree with results obtained during fictive locomotion in neonatal rodents where 5HT_1A_ was reported to have an inhibitory effect on the spinal rhythmic activity (Beato and Nistri,1998, Dunbar et al.,2010, Liu and Jordan,2005, Pearlstein et al.,2005). For example, in the brainstem-spinal cord of neonatal mice, 5-HT release during fictive locomotion was enhanced by citalopram, a selective re-uptake inhibitor, and a decreased burst duration and amplitude was observed (Dunbar et al.,2010). Since selective 5-HT_1A_ and 5-HT_1B_ antagonists reversed the inhibitory effect of citalopram, it was concluded that these receptors may rather be involved in rhythm inhibition. A similar conclusion has been drawn with neonatal rats where blocking 5-HT_1A/1B_ receptors during motor activity, produced by brainstem stimulation, induced speed-up of the rhythm (Liu and Jordan,2005). In both these studies, locomotor speed was impaired by 5-HT_1A_ receptor but the alternate pattern of locomotor rhythm was not blocked.

This discrepancy between results obtained during fictive locomotion or locomotion over a treadmill, when there is some exteroceptive stimulation, suggest that sensorimotor control is fundamental to the pro-locomotor effect of buspirone. Many studies employing different methodologies to induce locomotion have disclosed that reflex modulation is associated with locomotor expression (Frigon et al.,2012, Grillner and Shik,1973, McCrea,2001). Similarly, buspirone treatment induces spinal reflex re-organization and promotes locomotor activity.

### Conclusion

In summary, even if the role of 5-HT on motoneuron excitability has been extensively studied for more than fifty years, our knowledge is still scarce on how this neuromodulator contribute to sensorimotor control. The heterogeneity of 5-HT receptors locations (pre-, intra- or extra-synaptically) make it really difficult to assess the outcome of a treatment with this neuromodulator after a complete spinal cord injury. Reflecting this heterogeneity, buspirone, if given at a dose that can trigger locomotion, was shown to have biphasic consequence on the H-reflex in time after an acute lesion of the spinal cord, starting with an early and acute inhibition, followed by an excitation of the reflex.

## AUTHOR CONTRIBUTIONS

Y.D.: designed the study, conducted the experiments, acquired data, analysed data, and wrote the manuscript. H.L.: conceived and designed the study; acquired data, analysed data, and wrote manuscript; obtained funding for this work. All authors have approved the final version of the manuscript and agree to be accountable for all aspects of the work. All persons designated as authors qualify for authorship, and all those who qualify for authorship are listed.

## ACKNOWLEDGEMENTS

Y.D. was supported by studentship from Université du Québec à Trois-Rivières (Department of Anatomy).

